# Leader cells in collective chemotaxis: optimality and tradeoffs

**DOI:** 10.1101/642157

**Authors:** Austin Hopkins, Brian A. Camley

## Abstract

Clusters of cells can work together in order to follow a signal gradient, chemotaxing even when single cells do not. Cells in different regions of collectively migrating neural crest streams show different gene expression profiles, suggesting that cells may specialize to leader and follower roles. We use a minimal mathematical model to understand when this specialization is advantageous. In our model, leader cells sense the gradient with an accuracy that depends on the kinetics of ligand-receptor binding while follower cells follow the cluster’s direction with a finite error. Intuitively, specialization into leaders and followers should be optimal when a few cells have more information than the rest of the cluster, such as in the presence of a sharp transition in chemoattractant concentration. We do find this – but also find that high levels of specialization can be optimal in the opposite limit of very shallow gradients. We also predict that the best location for leaders may not be at the front of the cluster. In following leaders, clusters may have to choose between speed and flexibility. Clusters with only a few leaders can take orders of magnitude more time to reorient than all-leader clusters.

---

Eukaryotic cells commonly chemotax, moving in response to a chemical gradient, to locate wounds and move in a developing embryo. Clusters of cells often chemotax differently than single cells, cooperating to improve their sensing abilities [1–5]. In cooperating, cells may specialize, with leaders sensing the chemical gradient while others follow [2, 3, 6, 7]. We expect specialization to be most important if there is a large difference between the information different cells have about the gradient orientation, as in sharp transitions. This may occur when cell clusters follow a gradient that is “self-generated,” i.e. when cells near the rear degrade or sequester chemoattractant [7–14], allowing a cell cluster to migrate over distances much longer than its size during development [7, 10, 13] and cancer metastasis [11, 14].

We develop a minimal model of cluster chemotaxis with leaders and followers. Specializing improves chemotactic velocity in both sharp transitions and near-linear gradients of chemoattractant, but is not always beneficial. It is also not always best for cells in the front of the cluster to lead—cells near the middle or back of the cluster can have more information. Specialization not only impacts migration speed, but also strongly increases the cluster’s reorientation time.

## Model

We parameterize the chemoattractant profile:

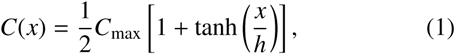

This function interpolates between step-like gradients and near-linear gradients depending on *h*, the scale of the transition from 0 to *C*_max_ (Fig. 1). We measure lengths in units of the cell diameter, so *h* = 1 is nearly steplike on the scale of a cell cluster. While the cluster moves in the *xy* plane, the position *x* in Eq. 1 is measured *relative to the lead cell at x* = 0– the cluster does not move relative to the gradient even as it moves in the lab frame. This is consistent with measurements of Sdf1 gradients in the zebrafish lateral line, which reach a steady-state in which they maintain their shape and move with the cluster [7, 10].

**FIG. 1.**
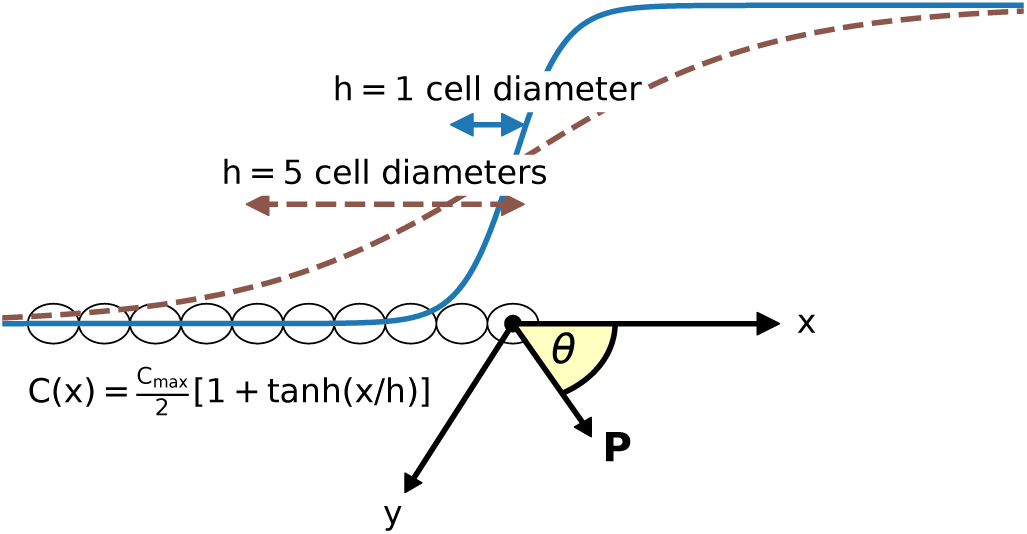
Geometry. The cell cluster is a rigid train with its front defining the point *x* = 0. The cells move in the *xy* plane, with polarity **P** = (cos *θ,* sin *θ*). *h* sets the width of the transition regime of the gradient in units of the cell diameter.

Leader cells make a measurement of the chemoattractant gradient direction. Earlier theory [15–20] and experiments [21–23] have established that a single cell sensing a chemical gradient is often limited in accuracy by the stochasticity of ligand-receptor binding [21]. We extend the model of [16, 17], assuming that leaders measuring the chemoattractant orientation have an angular error limited by ligand-receptor interactions. In the shallow gradient limit, this error 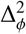 is [16, 17]

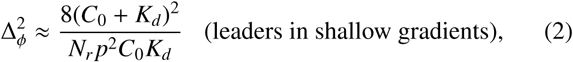

where *C*_0_ = *C*(*x*) is the mean concentration near the cell, *K*_*d*_ is the ligand-receptor dissociation constant, *N*_*r*_ is the number of receptors, and 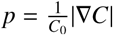 is the percent change in concentration across the cell. This shallow-gradient assumption may fail at sharper transitions (e.g. *h* = 1), so we determine the uncertainty 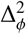 without this approximation by numerical integration [24]. We plot the leader angular error Δ_*l*_ as a function of the position within the cluster in Fig. 2. The follower uncertainty Δ_*f*_ is independent of position – i.e. we assume this noise arises from a process independent of chemosensing.

**FIG. 2.**
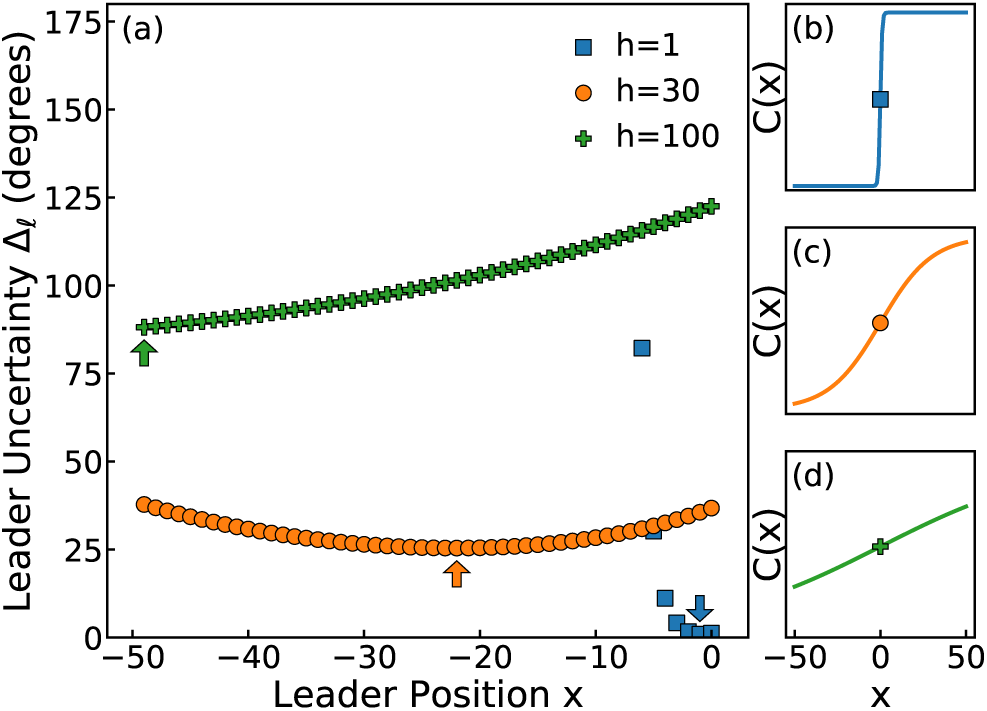
Directional Uncertainty of Leaders. (a) The uncertainty Δ_*φ*_ in a leader’s measurement of the gradient direction as a function of leader position *x* for sharp (*h* = 1), intermediate (*h* = 30), and shallow (*h* = 100) gradients. Arrows indicate the minimum of the curves, showing where the first leader will be added in a 50-cell train. (b)–(d) show *C*(*x*) corresponding to *h* = 1, *h* = 30, and *h* = 100, respectively. *C*_max_ = 2*K*_*d*_ here and throughout the paper.

To relate uncertainties in sensing to cell motion, we describe cells as actively moving and reorienting particles. Each cell *i* has an orientation *θ*_*i*_, corresponding to the cell being polarized in the direction **P**_*i*_ = (cos *θ*_*i*_, sin *θ*_*i*_). Here, leader cells align to the chemoattractant direction 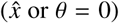, while follower cells follow the cluster’s direction *θ*_*c*_ (Eq. 5):

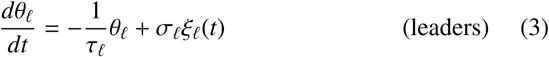

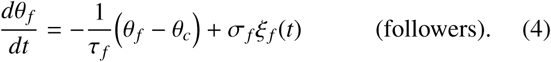

*σ* and *τ* for the leaders and followers depend on their accuracies Δ_*l, f*_, and may vary depending on cell position. *ξ*_*i*_(*t*) is Gaussian white noise with 〈*ξ*_*i*_(*t*)〈 = 0, 〈*ξ*_*i*_(*t*)*ξ* _*j*_(*t*′)〉 = *δ*(*t* −*t*′)*δ*_*i j*_. Angles *θ*_*ℓ*_ and *θ* _*f*_−*θ*_*c*_ are interpreted modulo 2*π*; we simulate Equations 3 and 4 by the Euler-Maruyama method with Δ*t* = 0.01 [24].

Collective migration is induced by having follower cells align to the cluster velocity direction *θ*_*c*_

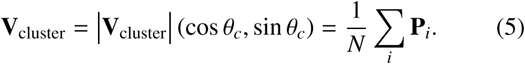

The cluster center of mass velocity is 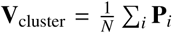 when cells are mechanically linked, and cell *i* would travel with velocity **P**_*i*_ in the absence of mechanical linkage [25]. The alignment of follower cells to the cluster orientation (Eq. 4) is a variant of the “self-alignment” [26] mechanism of Szabo et al. [27], and others [28–30] who showed that when cells align their polarity with their velocity, mechanical interactions between cells cause cells to align and migrate collectively. Our follower model is precisely that of [27] if cell velocity and cluster velocity are equal, i.e. the cluster is rigid.

We choose a dependence *σ*(Δ) and *τ*(Δ) so that the correlation time ***T*_P_** of a single cell’s polarity **P**_*i*_ is equal to 1 at all Δ [24]. This corresponds to the assumption that at all levels of uncertainty, a cell reorients at the same timescale, which we choose as our unit time.

## Optimal leadership strategies depend on chemoattractant profile and follower accuracy

A cluster may, depending on the chemoattractant profile *C*(*x*) and the accuracy of its followers Δ_*f*_, improve its mean velocity in the chemoattractant direction 〈*V*_*x*_〉 by specializing to leader and follower roles (Fig. 3) [31]. This is similar to results from a more complex model [32]. We do not assume a specific mechanism that determines which cells lead and which ones follow. Rather, we explore how a cluster behaves as the number of leaders changes. Here, and elsewhere, we add leaders from most to least accurate. We initially study a linear train of cells (Fig. 1), both for simplicity and as the most relevant geometry for narrow, extended systems like the zebrafish lateral line [33].

**FIG. 3.**
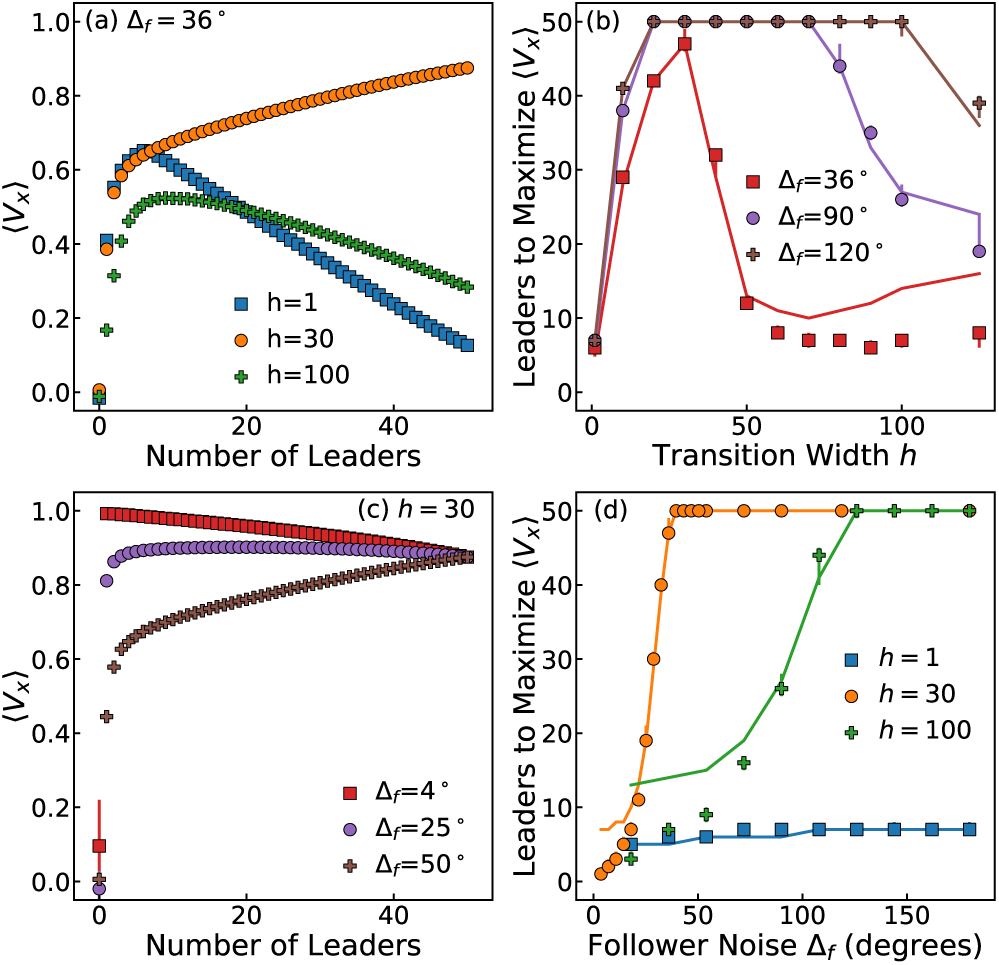
Cluster velocity depends on leader number. Trains of *N* = 50 cells. Symbols are simulations with 95% confidence intervals from bootstrapping; lines are the independent follower approximation [24]. (a) For sharp (*h* ∼ 1) and wide (*h* ∼ 100) gradients, the cluster can migrate substantially more quickly in the gradient direction with fewer leaders. (b) The number of leaders that maximizes 〈 *V*_*x*_〉, 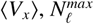, increases, then decreases as a function of gradient transition width *h*. (c) At fixed *h* = 30, raising the follower noise Δ _*f*_ lowers 〈 *V*_*x*_〉 at leader fractions less than one. (d) 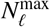 transitions from low to high for medium and wide gradients, but remains relatively constant for sharp gradients as the follower noise is increased.

For sharp transitions (*h* = 1), 〈*V*_*x*_〉 first increases and then decreases as we increase the number of leaders, reaching a maximum at 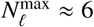(Fig. 3a). Consistent with our intuition that specialization will be most effective when transitions are sharp, 〈*V*_*x*_〉 increases monotonically in the number of leaders for the wider transition (*h* = 30). By contrast, in the nearly-flat concentration profile of *h* = 100, chemotactic velocity is maximized by having only a few cells be leaders (Fig. 3a).

To better understand how the optimal number of leaders depends on the chemoattractant profile, we study 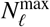, the number of leaders which maximizes 〈*V*_*x*_〉 (Fig. 3b). 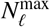 is small at sharp gradients, and initially increases as the transition size *h* increases – reflecting that for sharp transitions, leaders even a few cells away from the transition have extremely high levels of uncertainty, and will not increase *V*_*x*_. We would expect that further increasing *h* places more cells in the transition region and would increase 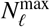 monotonically. Instead, we see that 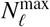 decreases at large *h* as the profile *C*(*x*) becomes nearly linear.

This apparently counter-intuitive result can be understood directly from the leader uncertainty Δ_*ℓ*_ as a function of cell position (Fig. 2). For gradients with a sharp transition regime (*h* = 1), leader uncertainty steeply increases for cells further away from the train front. At wider gradients (*h* = 30), there is a smaller difference between the best and worst leader. In fact, the most effective leaders tend to be in the middle of the cluster. As the gradient becomes near-linear (*h* = 100), instead of having cells with equal levels of uncertainty, cells near the back of the cluster have significantly lower uncertainty. This is because in linear gradients, the percentage change across the cell *p* is maximized farther from the transition, where the baseline concentration is lower, and *p* limits accuracy (Eq. 2). Specialization is rewarded at large *h because* there is a relevant difference in information available across the cluster.

Specialization relies on followers accurately using information from the leaders; the follower noise Δ _*f*_ can qualitatively change how 〈*V*_*x*_〉 depends on the number of leaders (Fig. 3c). If follower noise is very low (Δ _*f*_ = 4°), one leader can guide the cluster more effectively than when all the cells are leaders. For larger follower noises, (*V*_*x*_) is maximized when every cell is a leader (Δ_*f*_ = 50°, Fig. 3c). This leads to an even more dramatic change in 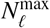: at *h* = 30, there is a rapid switch from 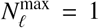 to 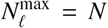 (Fig. 3d) as the follower noise is increased. However, this switching depends on the gradient width *h*. For the sharp *h* = 1 profile, 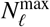 does not change much as Δ_*f*_ increases – having a small number of leaders is a *robust* strategy in sharp transitions. In the wider *h* = 30 and *h* = 100 gradients, increasing the follower noise level causes the number of leaders which maximizes 〈*V*_*x*_〉 to switch from low to high. Again, this can be understood by referring to Fig. 2. When there is a large difference between the best and worst sensors (*h* = 1), the magnitude of follower noise is relatively unimportant: Δ_*f*_ is usually larger than Δ_*ℓ*_ for the few well-informed cells, but smaller than Δ_*ℓ*_ for the bulk of the cells. By contrast, for *h* = 30, most cells have roughly the same amount of information about the gradient direction, and changing Δ _*f*_ can rapidly switch between Δ _*f*_ > Δ_*ℓ*_ for all cells, in which case it is optimal to have all cells be leaders, and Δ _*f*_ *<* Δ_*ℓ*_ for all cells, when as few cells as possible should lead.

Though our model of Eqs. 3-4 has a complex long-range collective interaction, we can quantitatively understand Fig. 3 with a much simpler independent follower model (solid lines in Fig. 3) [24]. Our independent follower model assumes that the follower error *θ*_rel_ ≡ *θ*_*f*_ - *θ*_*c*_ is independent of *θ*_*c*_, and also assumes *θ*_*c*_ ≈ *θ*_*L*_, where *θ*_*L*_ is the angle of only the leaders,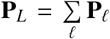. Effectively, each follower then independently follows the leader cells. The independent follower approximation is most effective at high levels of follower noise Δ _*f*_ (where follower-follower correlations are shorter-lived and less important), and sharp gradients (low *h*).

## Cluster reorientation and leader strategy

Because of the correlations between followers, a collectively sensing cluster can be highly persistent – even if it is moving in an incorrect direction. To understand this persistence, and the time it takes to reorient, we compute the velocity autocorrelation function 〈*δ***V**(*t*) *δ***V**(*t*′)〉 and fit it to an exponential to find the cluster’s correlation time *T*_*c*_ [24]. A short correlation time could be advantageous if cell clusters need to rapidly change direction (e.g. metastasizing clusters [34]), while long correlation times could be preferred for cell clusters traveling in consistent directions that must resist perturbations in the concentration profile (e.g. the zebrafish primordium). Experimentally, larger cell clusters exhibit slower reorientation in electric fields [35] and generally slower spontaneous reorientation in confinement [36], so we study both the effect of the cluster size and the number of leaders.

Cluster velocity and correlation time vary significantly with cluster size *N* and the number of leaders (Fig. 4). For sharp gradients (*h* = 1), specialization *N*_*ℓ*_ *< N* improves 〈*V*_*x*_〉for all *N* >5; chemotactic velocities decrease sharply for the all-leader (*N*_*ℓ*_ = *N*) case at larger cluster sizes, as more and more uninformed cells are leading. Most strikingly, in larger clusters, as the number of leaders is decreased from *N*_*ℓ*_ = *N*, the correlation time *T*_*c*_ increases over two orders of magnitude (Fig. 4b). These changes are reminiscent of those observed in a much more complex model of [37]. *T*_*c*_ also increases with smaller leader numbers at large *h* [24].

**FIG. 4.**
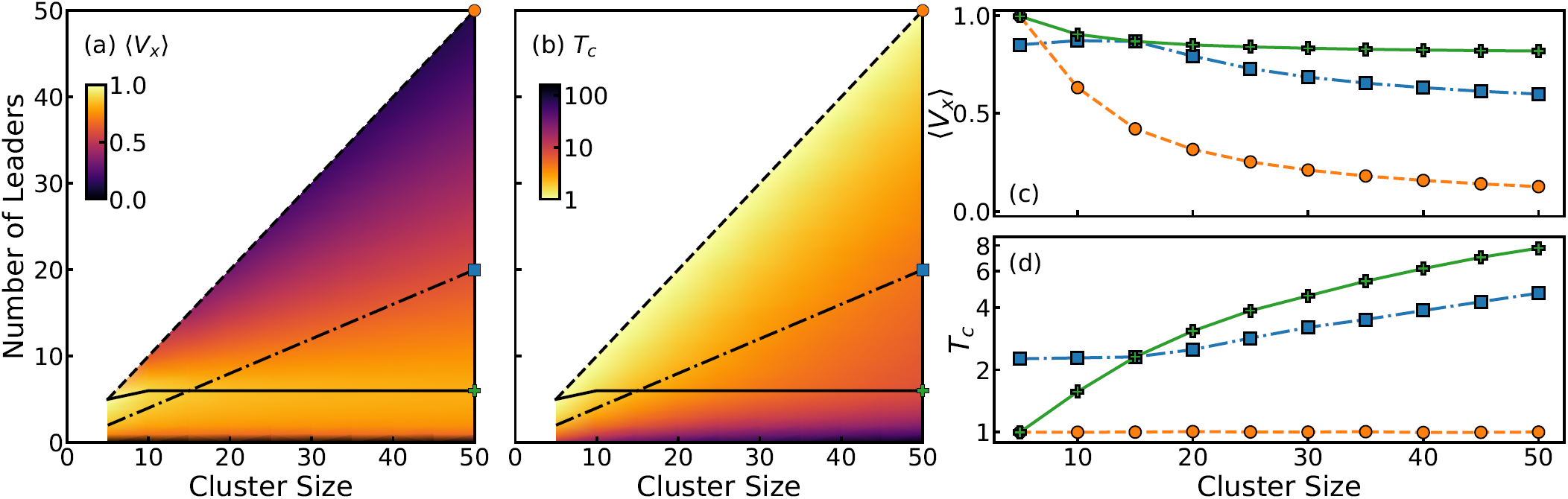
Cluster size and leader strategy alter the mean velocity and correlation time. Trains of *N* = 50 cells in sharp *h* = 1 gradient with Δ _*f*_ = 36°. Lines represent strategies of allocating leaders (not the independent follower approximation as in Fig. 3). The solid line (green pluses) corresponds to the number of leaders which maximizes *V*_*x*_. The dashed line (orange circles) corresponds to a leader fractions of 1 and the dot-dashed line (blue squares) corresponds to a leader fraction of 2/5. (a) The mean velocity in the gradient direction *V*_*x*_ as a function of cluster size and number of leaders. The corresponding slices are plotted in (c). (b) The cluster correlation time *T*_*c*_ as a function of cluster size and number of leaders. At fixed cluster size, this time can vary over two orders of magnitude from low to high leader number. The corresponding slices are plotted in (d). 95 % confidence intervals from bootstrapping in (c) and (d) are smaller than the symbol sizes.

Experiments often track gradient-sensing responses as a function of cluster size [1, 3, 35, 38, 39]. Our results show that, without specifying how leaders are chosen, even the qualitative dependence of chemotactic velocity or correlation time on cluster size are not known. We show how 〈*V*_*x*_〉 and *T*_*c*_ depend on cluster size with three leader allocation strategies (100% leaders, 40% leaders, and choosing the number of leaders that maximizes 〈*V*_*x*_〉) in Fig. 4cd. When all cells are leaders, 〈*V*_*x*_〉 monotonically decreases while *T*_*c*_ remains constant. If a fixed fraction 2*/*5 of cells are leaders, 〈*V*_*x*_〉 first increases, then decreases in cluster size, while *T*_*c*_ increases. And if we choose the leaders to maximize 〈*V*_*x*_〉, we see that larger clusters do not slow much – but they do see a steep increase in *T*_*c*_. These effects depend on the gradient shape, and the situation is much different for wider gradients, where increasing cluster size can increase 〈*V*_*x*_〉or even decrease *T*_*c*_ [24].

Though our results so far are for linear clusters, the most critical qualitative features (e.g. Fig. 3c) are consistent between compact and linear clusters [24]. The deviations between compact and extended clusters are easily understood in terms of the curves for Δ_*ℓ*_. For instance, in sharp transitions (*h* = 1), compact clusters have a higher 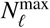, because there are more cells close to the transition *x* = 0.

We have also assumed that the only source of leader error in gradient sensing is ligand-receptor binding. Intracellular noise may also be significant at sharp enough gradients [21]. We study adding a downstream intracellular noise in quadrature,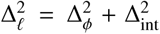. The qualitative dependence of 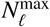 does not significantly change for low (∼4°) or moderate (∼18°) levels of downstream noise, but becomes substantially washed out at high (∼36°) levels [24]. This is expected: if the primary source of measurement error does not depend on the environment, specialization to leaders and followers will not be environmentally-dependent.

Our results show that choosing whether to specialize to leader and follower cells in a chemotaxing cluster is subtle. Clusters can chemotax more quickly if some cells sense and some cells follow, but the number of leaders to maximize directed migration depends heavily on the gradient and the accuracy of follower cells. In addition, cells at the front of the cluster may not be the most informed. Cells near the middle or back may be provide more accurate directional cues, consistent with experiments indicating that collectives are not necessarily steered by the cells at their front [40]. Our work shows rear-steering might be optimal in shallow gradients. Cluster persistence times are dramatically increased by leader specialization, and different strategies of allocating leaders trade off chemotactic speed with a cluster’s ability to reorient.

Are chemosensitive cells within clusters positioned according to our model? Experiments on the zebrafish lateral line primordium demonstrated that a small fraction of chemosensing cells at the front can restore the migration of a cluster when the rest of the cells have reduced chemosensing ability [41], suggesting that small numbers of sensing cells at the front can guide a cluster – consistent with our results. The scale of the transition from low to high concentration is set by the ratio between *D*, the effective diffusion coefficient of the ligand, and the degradation rate *k_* − [10]. [10] found *D* = 5 µm^2^/s for Sdf1 and *k*___ = 0.0003s^-1^ led to good agreement with their data, giving 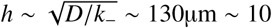 cell diameters [42]—consistent with leaders near the front of the cluster. Future experiments determining how varying the chemosensing ability of cells at different positions within a cluster will be an important way to test the leader-follower mechanism and whether the cells best able to sense the gradient are the ones that drive the cluster’s directionality.

We thank Holger Knaut for useful conversations, and Alma Plaza-Rodriguez for performing initial simulations on a related model. We acknowledge funding from an IDIES Seed Grant and Johns Hopkins University.

## Supplemental Information

### 1 Determining *σ* and *τ* as a function of angular noise Δ

We want Equations 3 and 4 to represent cells with given angular uncertainties Δ_*i*_. How can we map between Δ and the parameters of the model for each cell, *τ* and *σ*? To answer this question, we have to understand a bit more about the solutions of these equations, which are of the form

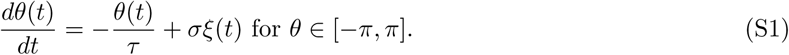

Without periodic boundary conditions, the dynamics of *θ* would follow an Ornstein-Uhlenbeck process [1], which can be described in terms of a known steady state distribution (normal with mean 0 and variance 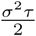, as in Equation S4) and a transition probability (Equation S6) which is the distribution of *θ* at a time *t*′ given the value of *θ* at an earlier time *t*. However, the periodic boundary conditions mean that the dynamics of *θ* are more complicated than a standard Ornstein-Uhlenbeck process. The steady state distribution becomes a truncated normal distribution determined from renormalizing Eq S4 on an interval [−*π, π*]. This results in the steady state probability distribution

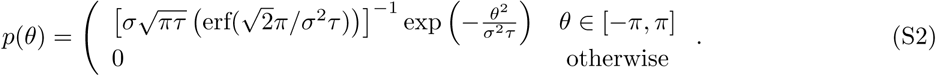

On the other hand, the transition probability is not analytically tractable. Therefore, we use results from the Ornstein-Uhlenbeck process in the limits of very small and very large noises Δ_*i*_, but must use numerical methods for intermediate values of Δ_*i*_.

For each cell *i*, the fluctuations of its angle *θ*_*i*_ about its mean value are characterized by the term 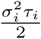 (Eq. S2). Therefore, we set the scale of these fluctuations equal to the angular noise Δ_*i*_ through the equation

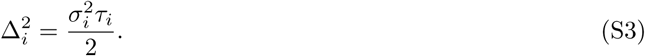

We note here that 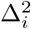 can be larger than 2*π* – here, we are setting the variance of the *parent normal* that is truncated to find Eq. S2.

If we applied only our formula for Δ_*i*_ (Eq. S3), there would not be a unique way to choose both *σ*_*i*_ and *τ*_*i*_ – we need to do more than just fix the variance of the angle 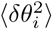. Therefore, we develop a procedure to choose values for *σ*_*i*_ and *τ*_*i*_ so that the behavior of the cell is realistic at all noise levels. If we naively chose *σ*_*i*_ ∼ Δ_*i*_, then as Δ_*i*_ became large, the cell would undergo angular diffusion with a diverging diffusion coefficient—physically unrealistic. We want to find functions *σ*(Δ) and *τ* (Δ) so that the correlation time *T***_P_** of a single cell’s polarity **P**_*i*_ is equal to 1 at all angular uncertainties. This corresponds to the idea that the cell has a constant time to reorient, independent of how accurately it is measuring its environment.

We first consider a Gaussian approximation, which does not account for the periodicity of the angle *θ*, to derive the asymptotic expressions for *σ*(Δ) and *τ* (Δ). Then, we numerically determine values for *σ*(Δ) and *τ* (Δ) at intermediate levels of angular uncertainties.

#### 1.1 Gaussian approximation

For an angle *θ*(*t*) relaxing to 0 with noise *σ* and time constant *τ,* the equation governing *θ* is Equation S1. However, in the Gaussian approximation, the periodicity of the variable *θ* is ignored. Although this result is not generally applicable, it is asymptotically correct in the limits of very small and very large angular noise. This is an Ornstein–Uhlenbeck process whose steady state distribution is a normal distribution with mean 0 and variance 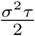

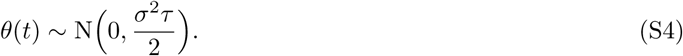

An angle at a later time *θ*(*t* + *t*′), given the value of *θ*(*t*), will have the distribution

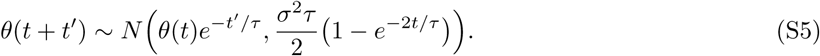

The first equation is the marginal distribution of *θ*(*t*) and the second equation is the conditional distribution of *θ*(*t* + *t*′) given *θ*(*t*), so the joint probability distribution *p* (*θ*(*t* + *t*′); *θ*(*t*)) is just the product of the two distributions. Let *θ*(*t*) = *x*_1_ and *θ*(*t* + *t*′) = *x*_2_ for ease of notation. Then, the joint probability density function is

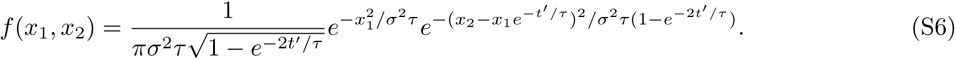

For a polarity vector **P**(*t*) = (cos*θ*(*t*),sin*θ*(*t*)), the time correlation function is given by

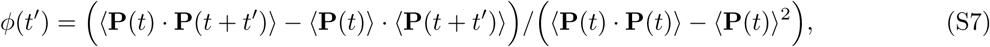

which in steady state only depends on the separation in time, and is normalized so that *ϕ*(0) = 1.

The term 〈**P**(*t*) · **P**(*t* + *t*′)〉 is given by the following expression:

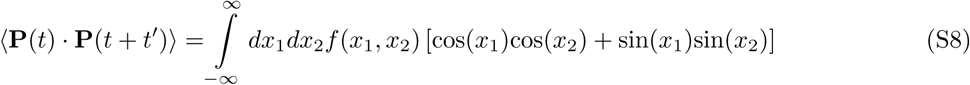

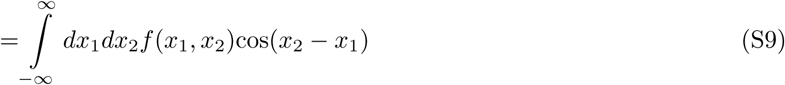

To evaluate the this term, we change the variables in the joint density function in Equation S6 through the following transformation:

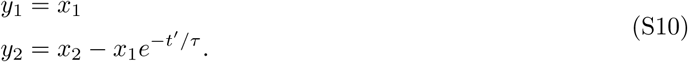

The Jacobian from this transformation is 1, so the new joint probability distribution is

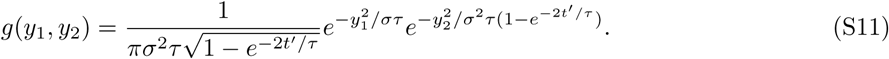

We can apply this transformation to Equation S9 and use the distribution in Equation S11 to evaluate the expectation value in the following way:

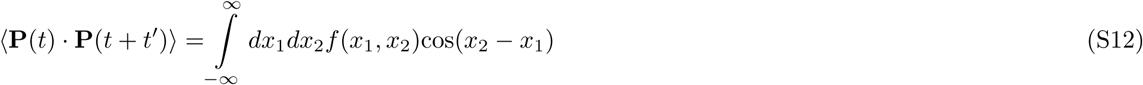

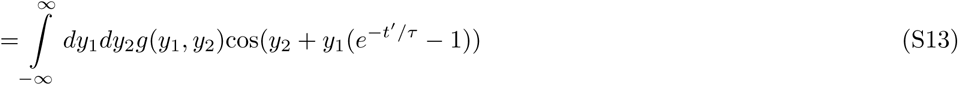

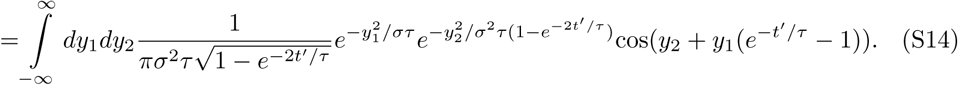

Applying the integral

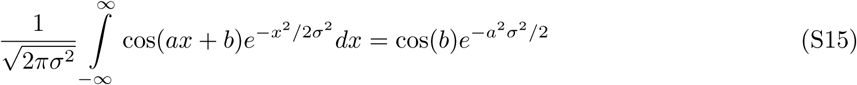

twice (once for *y*_1_ and once for *y*_2_) to Equation S14 gives the result

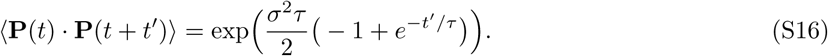

In steady state, the mean polarities will be independent of time, and the identity

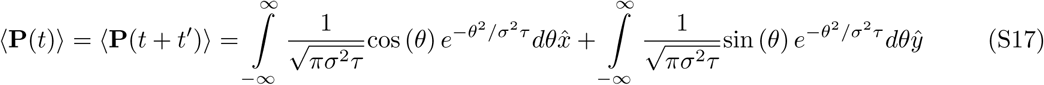

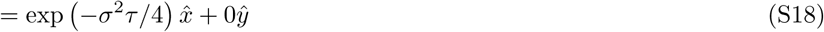

holds from applying Equation S15. Therefore, the term 〈**P**(*t*)〉 · 〈**P**(*t* + *t*′)〉 evaluates as

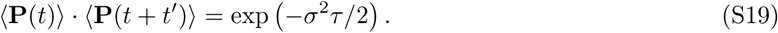

Thus, the final expression for *ϕ*(*t*′), normalized by its value at *t*′ = 0 is

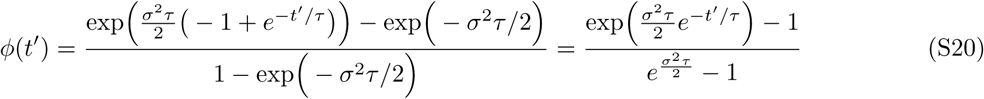

The correlation time is defined as the integral of this correlation function from *t*′ = 0 to *t*′ = ∞. Only the numerator of Equation S20 needs to be evaluated since the denominator does not depend on *t*′. Therefore, the integral to evaluate is the integral *I*:

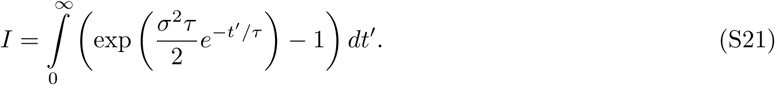

With the substitution 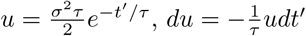, the expression for the integral becomes

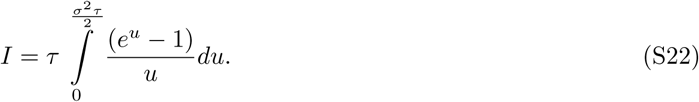

This integral can be solved and gives

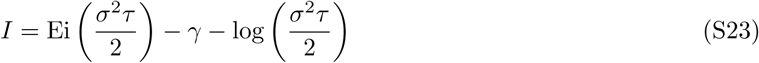

where *γ* is the Euler–Mascheroni constant and the exponential integral Ei is defined as

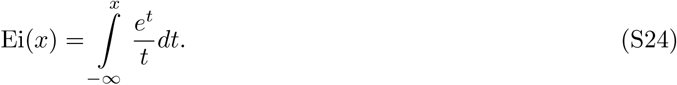

Thus, the correlation time, *T***_P_**, of the polarity **P** is

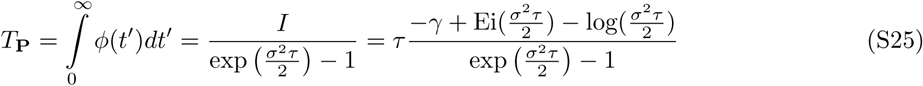

in terms of the angular relaxation time *τ* and angular noise *σ*.

To ensure that at every level of angular noise the polarity correlation time is constant, the polarity correlation time is set equal to 1 and the angular relaxation time *τ* and angular noise *σ* are chosen so that the angular uncertainty is as required. This can be described by the following equations

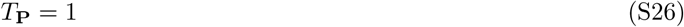

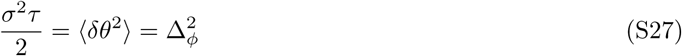

so that once a cell’s angular uncertainty 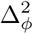 is known, the correct *τ* and *σ* for that cell can be chosen to set its polarity correlation time equal to 1. Thus, the formulas for *τ* (Δ_*φ*_) and *σ*(Δ_*φ*_)

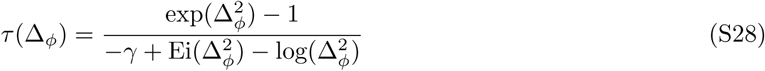

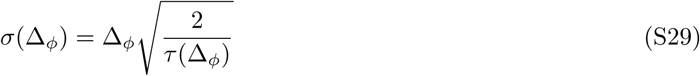

However, these formulas are only asymptotically correct because they do not account for periodic boundary conditions. Thus, the two limits we use give the following equations for the asymptotic forms

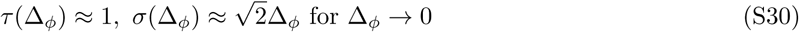

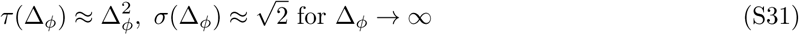

#### 1.2 Numerical Interpolation

For intermediate values of 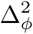 the parameters *τ* and *σ* determined from the Gaussian approximation can give real correlation times that deviate by up to 25% from the desired value of 1. This deviation is due to the wrapping of *θ* on the interval [-*π, π*], which is not accounted for in the Gaussian approximation. To accurately incorporate the effects of periodic boundary conditions, we want to numerically find values for the parameters *τ* (Δ_*φ*_) and *σ*(Δ_*φ*_) so that the relations *T***_P_** = 1 and 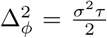 are both true. We simulate an angle following Equation S1. We ensure that *θ* remains on the interval [−*π, π*] by adding *π*, computing *θ* modulo 2*π*, then shifting *θ* back to the interval [-*π, π*] by subtracting *π*. That is, on each time step, we apply the following formula

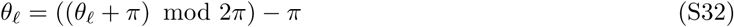

In the Gaussian approximation, the quantity 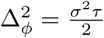 is the variance in the steady-state Gaussian distribution of *θ*. With periodic boundaries, the steady-state distribution of *θ* is a truncated normal distribution on [*-π, π*]. Therefore, 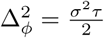 is the variance of the parent normal distribution to truncated normal distribution for *θ* (as in Equation S2). Thus, small values of 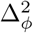 result in an approximately Gaussian distribution of *θ* around the gradient direction while very large values of 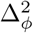 correspond to a uniformly distributed value of *θ*, corresponding to a cell that chooses its direction randomly. However, the transition probability from one angle to another is not analytically tractable with periodic boundaries, so we need simulations to determine the correlation time *T***_P_**.

As implied by the form of Equation S25 and the Buckingham Pi Theorem, the equation for the correlation time can be written in terms of the parameters *τ* and *σ* as

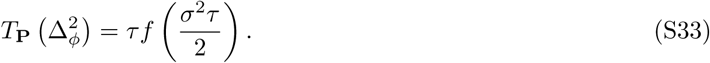

This is also shown empirically in Figure S1, where for various values of *τ* and 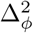 the polarity correlation time 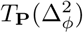 is determined from simulations with 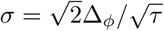. Therefore, without loss of generality, we choose the parameter *τ* = 1 and 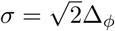 and simulate *T***_P_** (Δ_*φ*_) to find the function 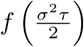. This function is useful because choosing 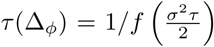 ensures that the correlation time of the polarity *T*_*p*_ = 1 at all values of Δ_**φ**_. Thus, we choose the parameters according to the rule

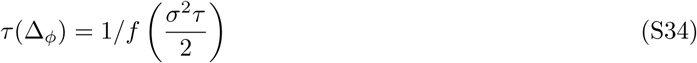

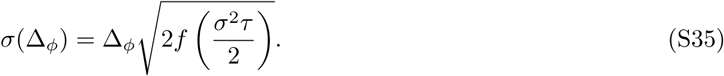

**Figure S1:**
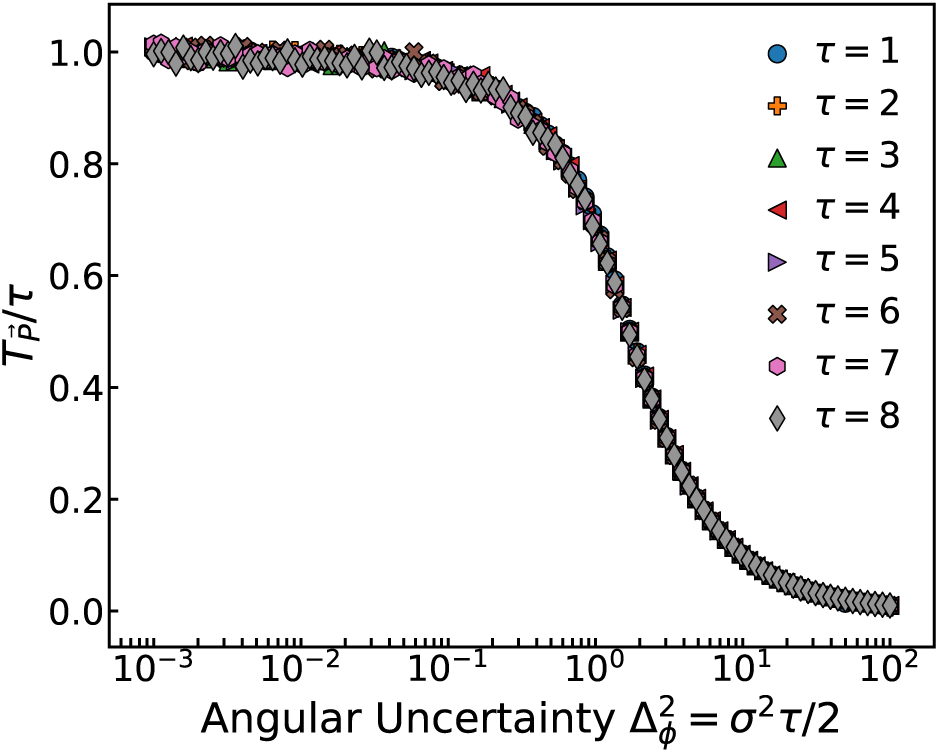
Universality of the single-cell polarity correlation time scaling. The scaling between the correlation time of the polarity 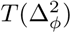 normalized by the parameter *τ* is universal for all values of *τ*. Here, *τ* is chosen as a parameter and 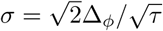

To find the correlation time, we simulate an angle relaxing to 0 with periodic boundaries enforced (see Simulation Details section below). We use a time step of Δ*t* = 0.01, as in the main simulations, and we simulate to a time of 3000, which was found to generate good statistics for the correlation function. The correlation time is found by fitting an exponential to the correlation function, though re-scaling the time at which correlation function reaches 1/2 to determine when a 1/e decay would have occurred gives similar trends. We repeat this procedure to generate 100 measurements of *T***_P_** for each value of 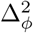. Then, the parameters *τ* (Δ_*φ*_) and are found through Equations S34 and S35 and linear interpolation. A grid is created for 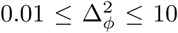 where, for each range of orders of magnitude, the grid spacing is 1/10 of the smallest value. Outside those limits, we use the asymptotic forms.

#### 1.3 Verifying Numerical Scheme

**Figure S2:**
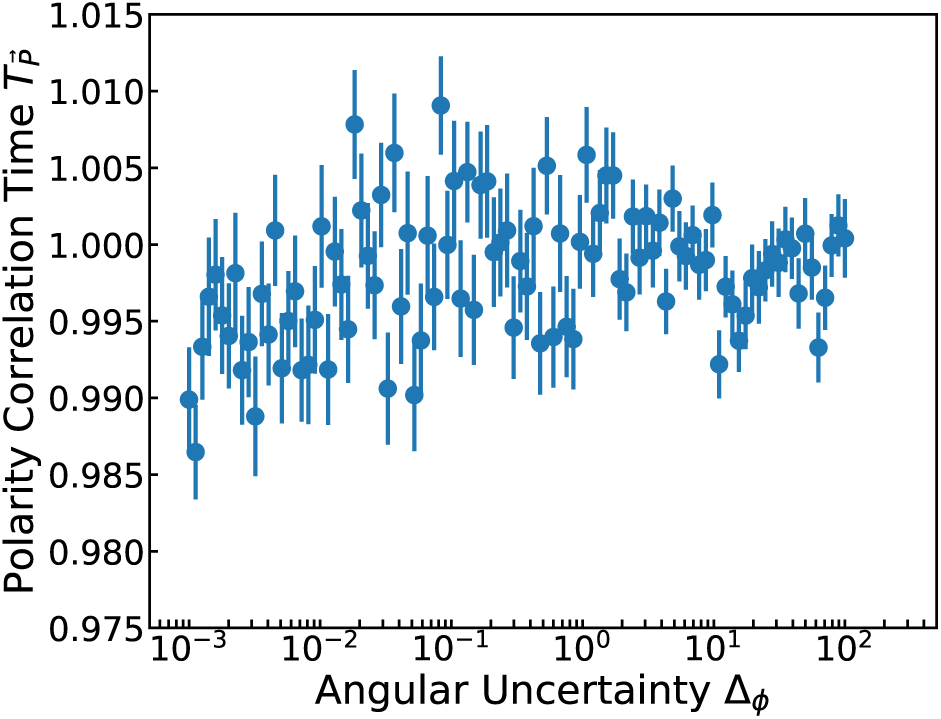
Simulated tests of the numerical scheme. The correlation time of the polarity *T***_P_** for a single cell relaxing to an angle of zero is plotted as a function of its uncertainty Δ_*ϕ*_. The parameters *σ*(Δ_*ϕ*_) and *τ* (Δ_*ϕ*_) are chosen according the process described above. The correlation time remains within two percent of the desired time of one.

To determine the accuracy of the numerical method, we compute *T***_P_** for single cells of various angular uncertainties Δ_*ϕ*_, where we choose the parameters *τ* and *σ* according to the procedure outlined in the previous section. In Figure S2, we present values computed for angular uncertainty levels that lie in both asymptotic limits and in the interpolation regime. The computed values are within at least 2% of the desired time of 1. Values outside the range considered here will be at least this close to 1 because the asymptotic limits will improve at very small or very large Δ_*ϕ*_.

### 2 Extending the Hu et al. result beyond the shallow gradient approximation

As in [2, 3], we consider a circular cell with *N*_*r*_ receptors uniformly spaced along its perimeter. Let *ϕ* denote the direction of the gradient. We model the receptors on the cell *x*_1_*…x*_*Nr*_ as *N*_*r*_ independent Bernoulli trials that can be occupied with value 1 or unoccupied with value 0. For simple ligand-receptor kinetics, the probability of receptor *n* being occupied given concentration *C*_*n*_ at the receptor and ligand-receptor dissociation constant 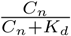 and the probability of being unoccupied is 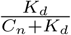. The probability that the *n*th receptor is occupied is a function of the gradient direction *ϕ*. Thus, the probability distribution function for the *n*th receptor is

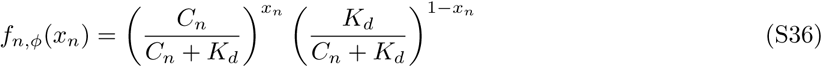

The likelihood function for a cell estimating the gradient direction *ϕ* given the values at the receptors is

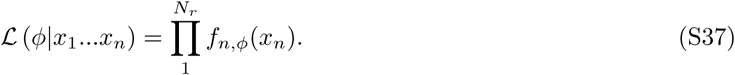

The log-likelihood function is

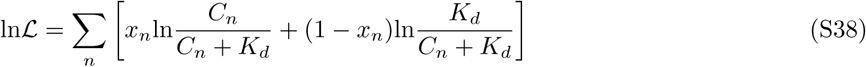

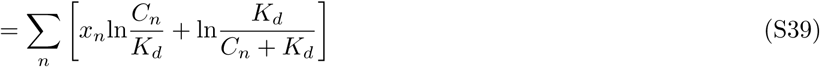

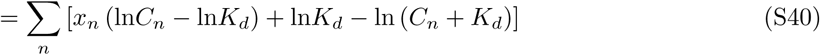

For an estimation of the gradient direction, we are interested in computing the second derivative of the log-likelihood function with respect to *ϕ*. Therefore, let a prime ′ denote a derivative taken with respect to *ϕ*. Then, the first derivative of the log-likelihood function with respect to *ϕ* is

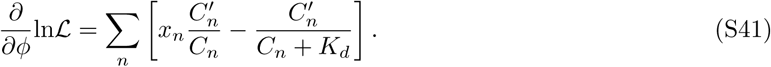

Taking another derivative gives

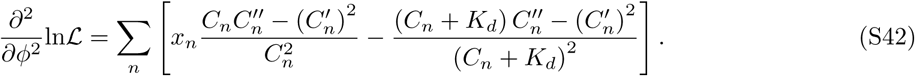

To find the expectation value, first note the for the Bernoulli trials *x*_*n*_, the expectation value is just the probability of occupancy 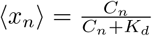. With this expression, the expectation value of the second derivative becomes

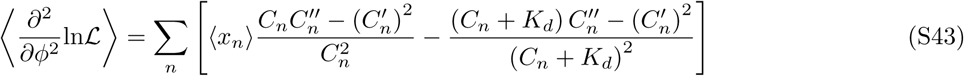

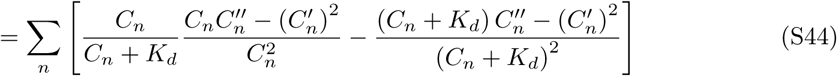

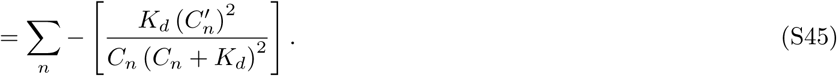

This expression holds true for any gradient profile and does not make any assumptions about its steepness.

In this work, the goal is to compute the uncertainty for leader cells with the gradient varying the in *x* direction given by 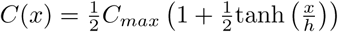. For the *i*th cell in the cluster, the *x* position of the center is 1-*i*, where *i* ranges from 1 to the cluster size *N*. Since we are working in units in which the cell diameter is 1, the *x* positions of the receptors are given by

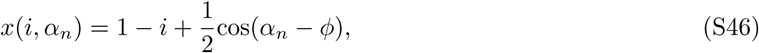

where *α*_*n*_ is the angular position of the *n*th receptor. The sum in Equation S45 can be approximated as an integral for a large number of receptors. Therefore, for a given cell *i*, the Fisher information is

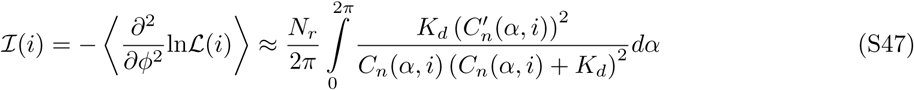

where the concentration and its derivative is written in terms of *α* and *i* using Equation S46. This expression does not depend on the gradient direction *ϕ* because the receptors are evenly spaced, and in the cell is symmetric under rotations in the continuum limit. We numerically integrate Equation S47 with Gaussian quadrature to determine the Fisher information for each cell in the cluster. With the Fisher information, the Cramér-Rao bound gives the minimum uncertainty for the gradient direction estimated by the *i*th cell as

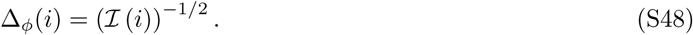

### 3 Independent Follower Approximation

The cluster velocity is given by the mean polarity of all the cells in the cluster, which can be decomposed into leader and follower contributions

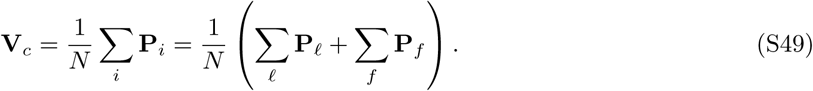

Therefore, the mean *x* velocity of the cluster is given by

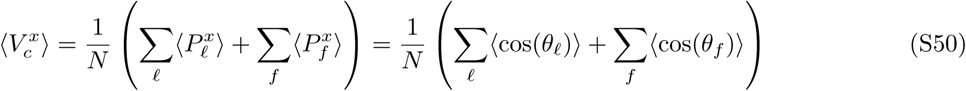

in terms of the leader angle *θ*_*ℓ*_ and follower angle *θ*_*f*_. The followers relax towards the angle of the cluster *θ*_*c*_ with some noise. The angle of a follower *θ*_*f*_ can be written in terms of the angle of the cluster and some angle *θ*_*rel,f*_, which is the angle of follower *f* relative to the cluster angle

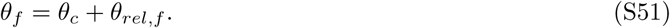

The *x* polarity of a follower is just the cosine of the follower angle, which can be rewritten using Equation S51 as

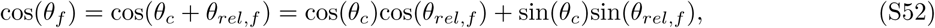

and the mean is given by

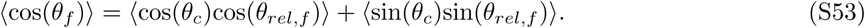

So far, the above results are exact. To develop the approximation, we make 2 key assumptions. First, we assume (1) that the relative angle of the follower *θ*_*rel,f*_ is independent of the cluster angle *θ*_*c*_, this can be simplified as

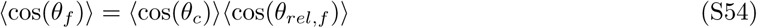

because both *θ*_*c*_ is symmetric about 0 so 〈sin(*θ*_*c*_)〉 = 0. To evaluate the expectation value 〈cos(*θ*_*rel,f*_)〉, we note that its dynamics are essentially that of the follower angle *θ*_*f*_ in a reference frame in which *θ*_*c*_ = 0. Therefore, the distribution of *θ*_*rel,f*_ will be a truncated normal distribution on [−*π, π*] with mean 0 and whose parent normal has a standard deviation Δ_*f*_. From this result, an exact expression for 〈cos(*θ*_*rel,f*_)〉 can be applied to find

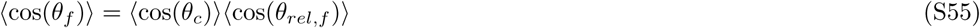

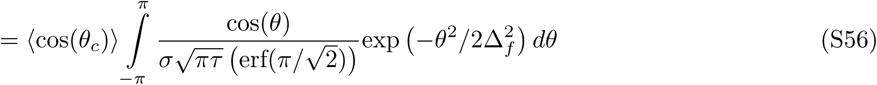

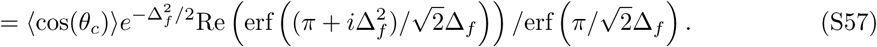

Let the contribution to the polarity from the leaders be defined as the total leader polarity vector **P**_*L*_

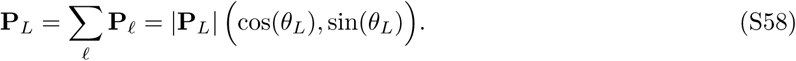

Then, we assume (2) that the average *x* component of the cluster is approximated by the average *x* component of **P**_*L*_, ie,

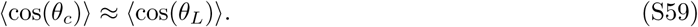

This assumption is generally good since the follower polarities align with the leaders in steady-state. However, it breaks down when the follower correlations contribute significantly to the accuracy of the cluster direction.

Explicitly, the term we use to approximate cos(*θ*_*c*_) is

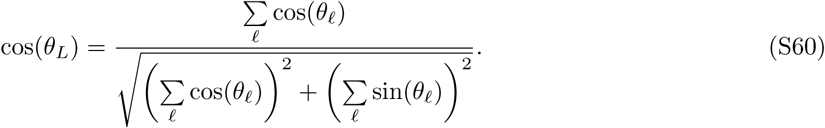

We compute 〈cos(*θ*_*L*_)〉 numerically. We draw the leader angles *θ*_*ℓ*_ from their respective steady-state truncated normal distributions and the compute the mean value of Equation S60 from the draws. We use 100,000 draws for *h* ≤ 100 and 500,000 draws for *h* = 125. Once this term has been computed numerically, Equation S57 gives the follower contribution 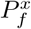 to the mean cluster velocity 〈*V*_*x*_〉, where we approximate the term 〈cos(*θ*_*c*_)〉 as 〈cos(*θ*_*L*_)〉. Then, the leader contribution 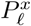 to 〈*V*_*x*_〉 is simply

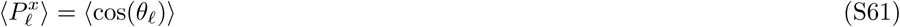

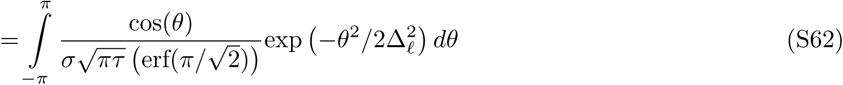

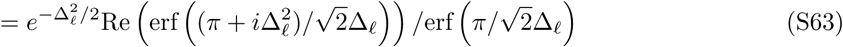

because the leaders independently align to the gradient direction. Thus, the final expression for the gradient velocity is

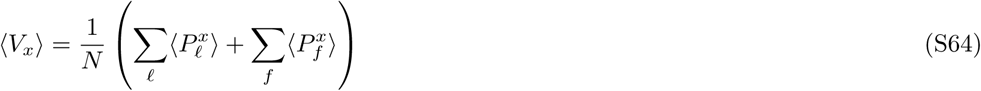

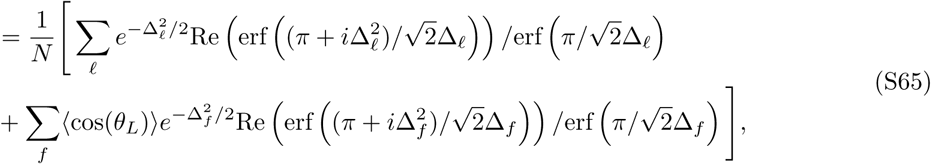

where 〈cos(*θ*_*L*_)〉 is computed numerically through S60, as described above. Once we have 〈*V*_*x*_〉 as a function of leader number, we can compute 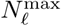.

To illustrate the signal that the followers respond to in this approximation, we plot cos (*θ*_*L*_) as leaders are added in Figure S3. This quantity does not vary as strongly as the *x* velocity because diffusing leaders do not distort the direction in a consistent way, which mitigates an inaccurate leader’s impact on the directionality of **P**_*L*_.

**Figure S3:**
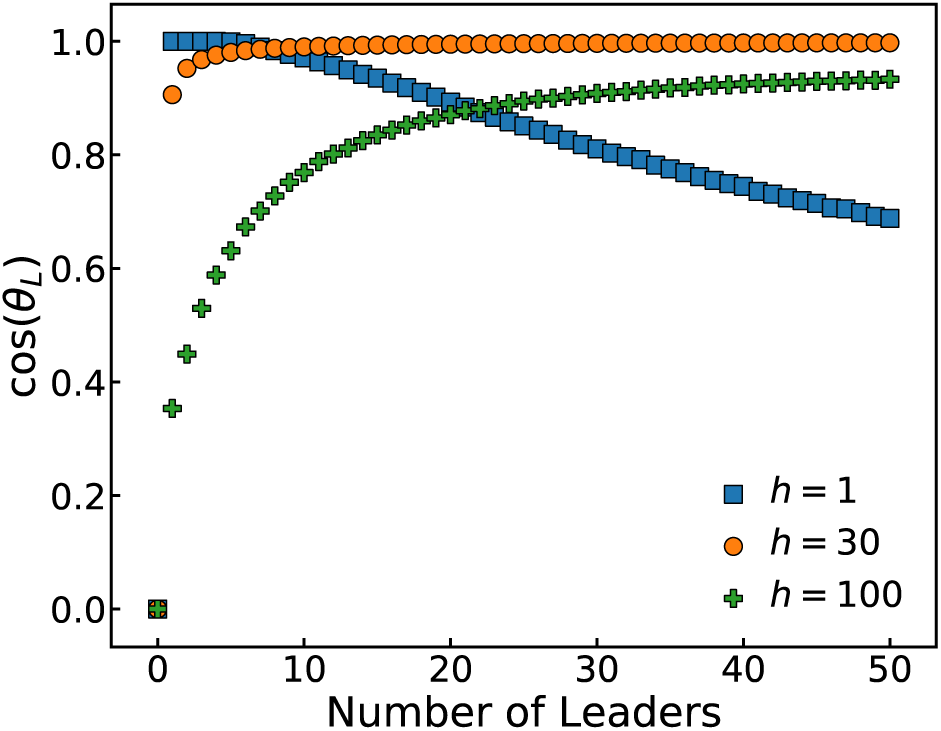
Leader signals in the independent follower approximation. In the independent follower approximation, the signal to the followers is the cosine of the polarity vector of all the leaders. For sharp gradients (*h* ∼ 1), the leaders are first accurate and the cosine is near 1, but it decreases as essentially random leaders are added. For intermediate gradients (*h* ∼ 30), the curve is flat and large as all the leaders have similar and low uncertainty in the gradient direction. For wide gradients (*h* ∼ 100), the direction of the leaders becomes more accurate as more are added, with diminishing returns on additional leaders.

### 4 Follower Noise and Correlation Time

In the main paper, we show that the cluster correlation time *T*_*c*_ depends strongly on the leader number and the cluster size. However, one might suspect that the correlation time might also depend on the follower noise level Δ_*f*_ because the followers maintain the cluster velocity at any instant, driving its persistence. Therefore, at a fixed gradient, we show the dependence of *T*_*c*_ on leader number for low (Δ_*f*_ = 36°), intermediate (Δ_*f*_ = 72°), and high (Δ_*f*_ = 120°) levels of follower noise in Figure S4. In the sharp (*h* = 1), medium (*h* = 30), and wide (*h* = 100) gradients, the trend is the same—increasing the follower noise decreases the correlation time at leader fractions less than one. Also, even at the highest level of follower noise considered here, there is still a significant change in *T*_*c*_ from a small leader number to the all leader case, and this effect persists at each gradient.

**Figure S4:**
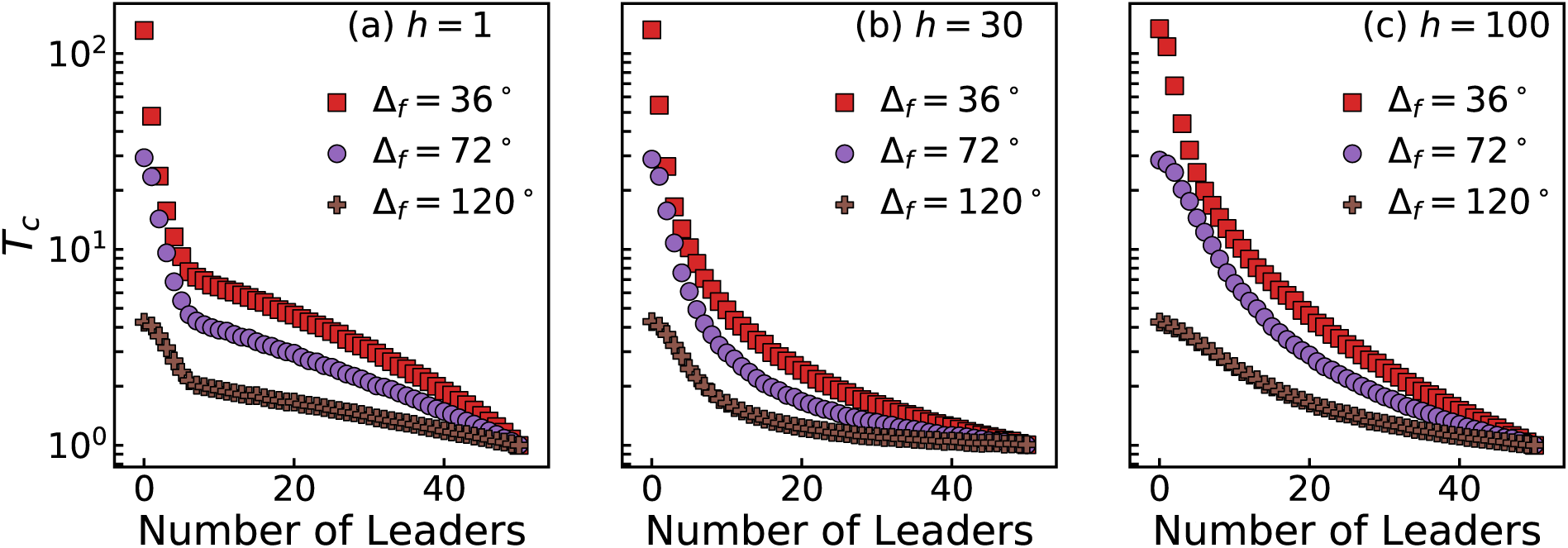
The cluster correlation time decreases with increasing follower noise. The cluster correlation time *T*_*c*_ decreases as a function of follower noise Δ_*f*_. Although the extent of the difference in cluster correlation time between the few leader and all leader case quantitatively depends on the follower noise, the trend does not. Shown are the gradients *h* = 1, *h* = 30, and *h* = 100 respectively in (a)-(c).

### 5 Cluster Geometry

We compare the train geometry considered in the main text to clusters with a compact geometry. We examine the geometry of a 4-layer oligomer (as considered in previous work [4, 5]), in which 61 cells are hexagonally packed in the cluster, as illustrated in Figure S5.

**Figure S5:**
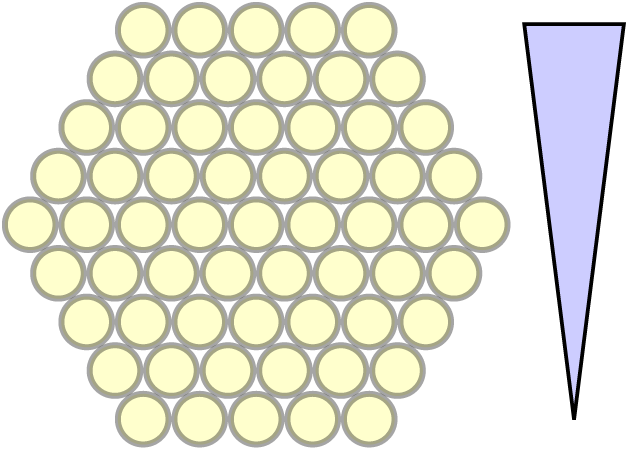
Illustration of a hexagonally packed cluster with 61 cells.

To exactly isolate the effects of geometry, we also simulate trains of *N* = 61 cells to compare with the more compact geometry. First, we examine how the geometry affects the mean migration speed, 〈*V*_*x*_〉. In Figure S6, we compare the mean velocity curves as a function of the leader number between trains and oligomers for small (*h* = 1), medium (*h* = 30), and large (*h* = 100) gradient widths at a fixed follower noise level Δ_*f*_ = 36°. For the sharp gradient, the hexagonally packed cluster has many more cells near the transition region, so it has many cells that can accurately measure the gradient. Therefore, adding leaders continues to increase its velocity for a larger number of leaders than the *N* = 61 train, and it can migrate more quickly than the train. In the medium gradient width, the accuracy of the leaders does not change significantly as a function of position. This is reflected in the very similar velocities between the geometries at all leader numbers in the *h* = 30 gradient. In the wide *h* = 100 gradient, the cells that can most accurately sense the gradient are near the back of each cluster. Therefore, the train migrates more quickly than the packed cluster, as the more extended geometry gives access to more accurate leaders. However, the differences between the geometries are not as striking as in the sharp gradient case because the *h* = 100 uncertainty curve does not vary as dramatically with position.

**Figure S6:**
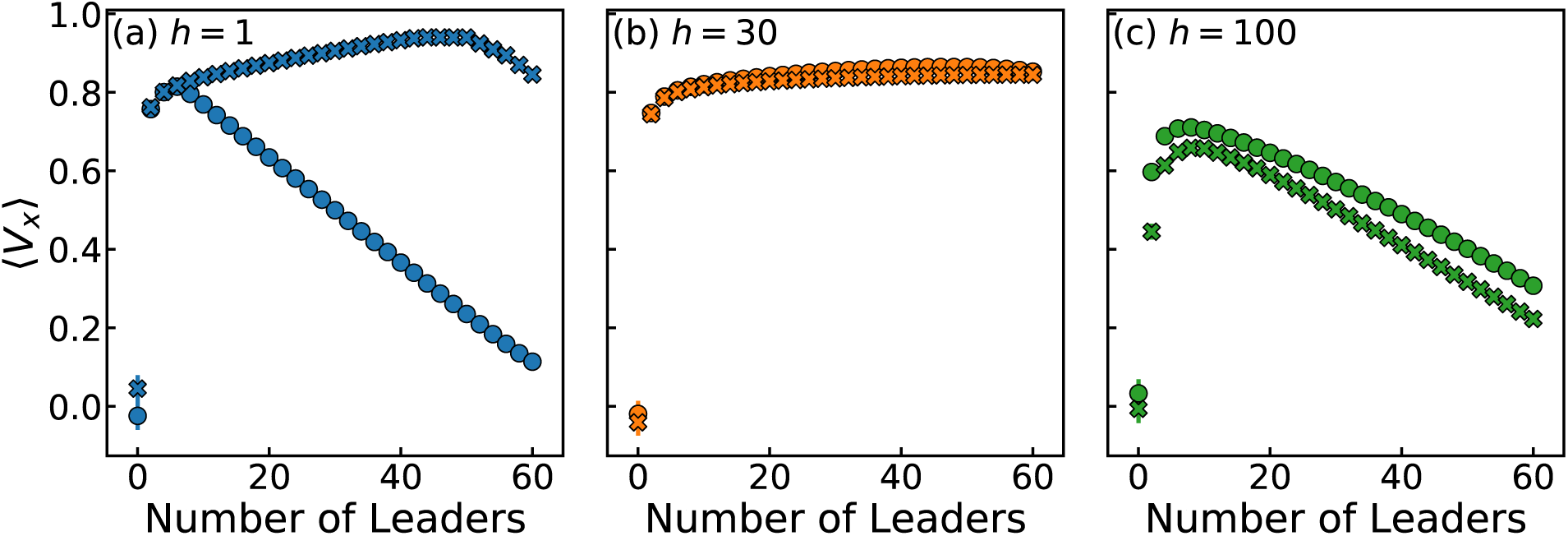
Cluster velocities for compact and extended clusters. We compare the mean velocity in the gradient direction 〈*V*_*x*_〉 between *N* = 61 trains and *Q* = 4 oligomers. The follower noise level is 36°. Circles represent the trains and Xs represent the oligomers. In the sharp *h* = 1 gradient of (a), the oligomer geometry places more cells near the sharp gradient transition, so there are more accurate leaders and the cluster can chemotax faster than the train. The *h* = 30 gradient in (b) produces a relatively flat leader uncertainty curve, so the geometry does not have a large impact. The wide *h* = 100 gradient in (c) means that the most information about the gradient direction is far from the transition regime, so the larger extent of the train geometry allows it to chemotax slightly faster than the oligomer.

One feature of the curves considered in Figure S6 is that the non-monotonic behavior of the velocity as a function of leader number is present in both geometries. We look to see if the non-monotonicity in the 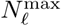 as a function of gradient width *h* also is robust to geometry. Figure S7 shows that for the *Q* = 4 layered oligomer 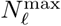 first increases, then decreases as a function of gradient width *h*. The hexagonally packed cluster starts out with a higher 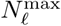 at low *h* because more of its cells are near the transition region. It also experiences a steeper initial decrease in 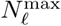 as *h* increases as the uncertainty curve changes from having a minimum near the transition regime to having a minimum far from the transition regime. This change more sharply affects the oligomer cluster because it does not extend as far in the *x* direction as the train. Finally, each has a similar number of leaders to maximize migration in the wide gradient limit. In this regime, the follower contribution becomes important relative to the leader signal. This is reflected in the breakdown of the independent follower approximation and the similar simulated values of 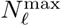 for both geometries.

**Figure S7:**
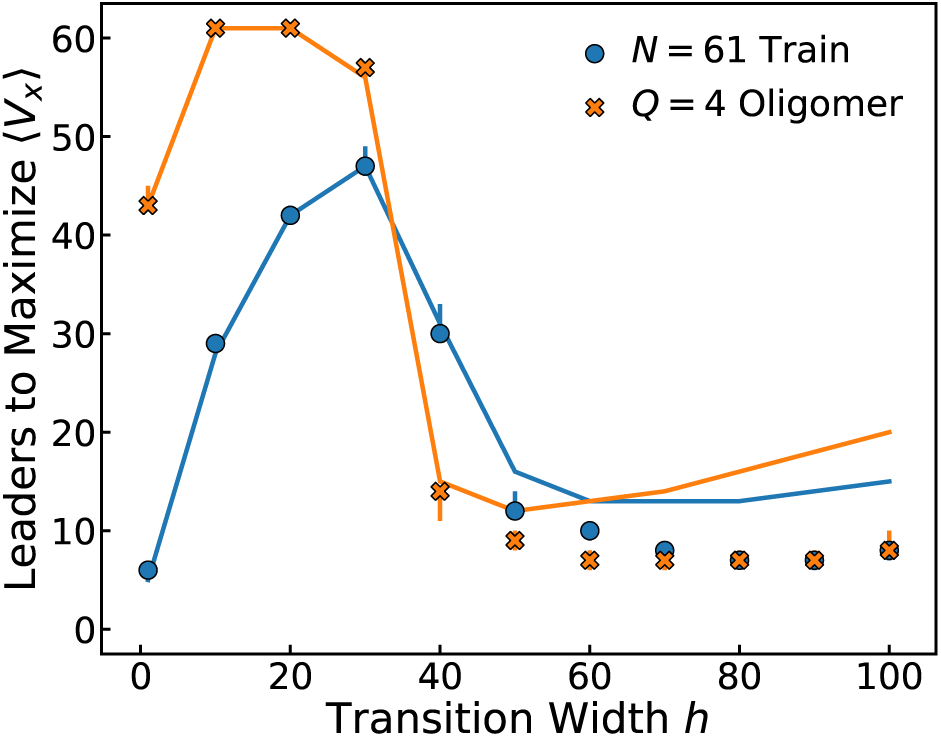
The number of leaders to maximize migration speed for compact and extended clusters. We compare 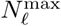 for *N* = 61 trains and *Q* = 4 oligomers. The follower noise level is 36°. Circles represent the trains and Xs represent the oligomers. Both geometries follow a similar trend - the number of leaders first increases then decreases as the gradient transition widens.

We also consider the cluster correlation time *T*_*c*_ for each geometry. The main qualitative trend that the correlation time can change dramatically from a small leader fraction to the all leader case, is robust to the geometry, as in Figure S8.

**Figure S8:**
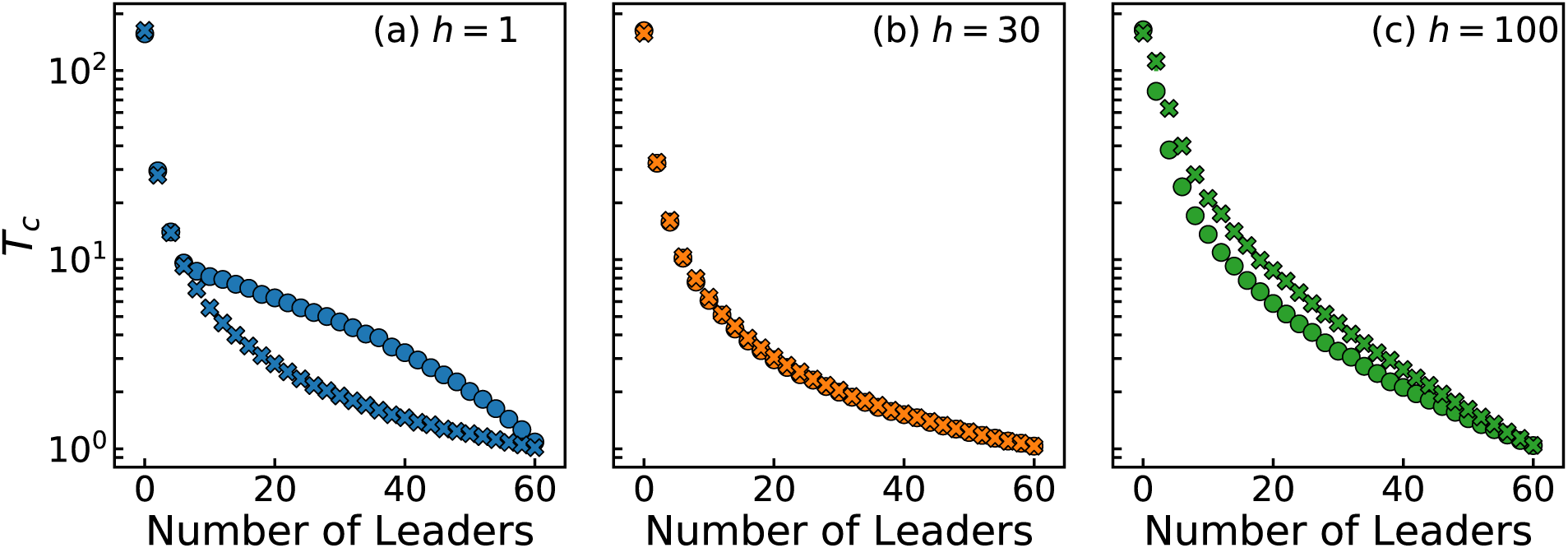
Cluster correlation time for compact and extended clusters. We compare the cluster correlation time *T*_*c*_ between *N* = 61 trains and *Q* = 4 oligomers. The follower noise level is 36°. Circles represent the trains and Xs represent the oligomers.

### 6 Intracellular Noise

In the main paper, we assume that the uncertainty in the leaders’ direction is due entirely to effects of the stochastic binding between ligands and receptors. However, additional noise may be introduced as the cell processes the information from the bound and unbound receptors. We vary the level of intracellular noise Δ_int_ that the leaders experience. We assume that this noise is independent of the ligand-receptor noise, and add it in quadrature with the directional uncertainty so that a leader’s uncertainty is 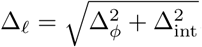. As the level of intrinsic noise increases, the effect of the gradient on leadership strategy washes out, as shown in Figure S9. However, the effects do not completely wash out until the level of intracellular noise approaches extremely high levels (∼ 72°).

**Figure S9:**
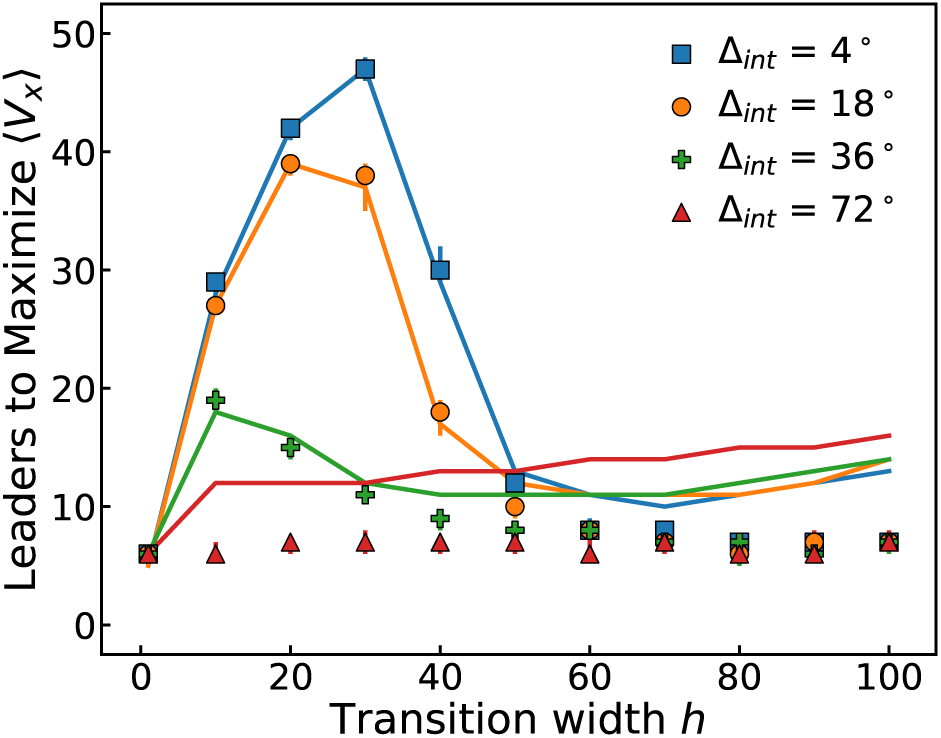
Qualitative features of leadership are robust to realistic levels of intracellular noise. Increasing the intracellular noise Δ_int_ eventually washes out the effects of the gradient on the number of leaders to maximize 〈*V*_*x*_〉, but for physically realistic values the qualitative features remain. The symbols are data for *N* = 50 trains with a follower noise of Δ_*f*_ = 36°, and the lines are the independent follower approximation predictions.

### 7 Additional Cluster Size Dependence Data

In the main text, we present data on how cluster size and leadership strategy affect the velocity and correlation time for a sharp gradient. Here, we consider the case of a wide *h* = 100 gradient. Unlike in the sharp gradient, the global maximum in velocity is achieved at the largest cluster size considered, *N* = 50, because larger clusters are more extended in the −*x* direction which gives them access to the cells that can most accurately sense the gradient.

**Figure S10:**
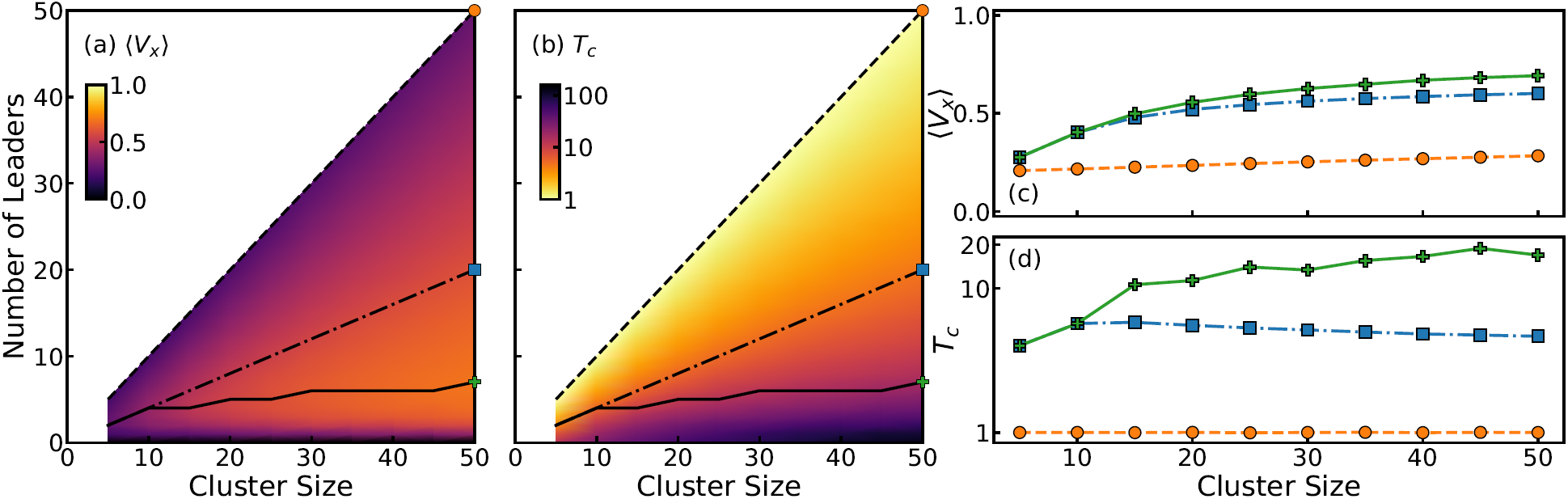
Cluster size and leader strategy in a wide gradient. Simulation data for a train of *N* = 50 cells in a wide *h* = 100 gradient with follower noise Δ_*f*_ = 36°. The solid line (green pluses) corresponds to the number of leaders which maximizes 〈*V*_*x*_〉. The dashed line (orange circles) corresponds to a leader fractions of 1 and the dot-dashed line (blue squares) corresponds to a leader fraction of 2/5. (a) The mean velocity in the gradient direction 〈*V*_*x*_〉 as a function of cluster size and number of leaders. The corresponding slices are plotted in (c), where the symbols are the data. (b) The cluster correlation time *T*_*c*_ as a function of cluster size and number of leaders. At fixed cluster size, this time can vary over two orders of magnitude from low to high leader number. The corresponding slices are plotted in (d). 95 % confidence intervals from bootstrapping in (c) and (d) are smaller than the symbol sizes..

### 8 Simulation Details

The cluster velocities are simulated until t=5,200 for various gradient widths, cluster sizes, follower noises, and number of leaders. Steady-state is considered to be reached after t=200, since that is approximately the longest correlation time encountered. From the steady-state data, we compute the correlation function *ϕ*(*t*′) = 〈**V**_*c*_(*t*) · **V**_*c*_(*t* + *t*′)〉− 〈**V**_*c*_(*t*)〉^2^ and determine a correlation time by fitting an exponential function to *ϕ*(*t*′)*/ϕ*(0). We show a representative example in Figure S11, which is the steady-state cluster velocity autocorrelation and its exponential fit for a train of *N* = 50 cells in an *h* = 10 gradient with 10 leader cells and followers with Δ_*f*_ = 36°. Re-scaling the time at which correlation function reaches 1/2 to determine when a 1/e decay would have occurred gives similar trends. After the correlation time *T*_*c*_ for the cluster has been measured for a run, the steady-state velocity data is broken up into intervals of 3*T*_*c*_ so that the mean of each interval is an independent measurement of the steady-state velocity. We repeat the simulation 50 times to generate 50 measurements of the correlation time and many measurements of the steady-state velocity. 95% confidence intervals are generated for the mean velocity in the gradient direction 〈*V*_*x*_〉 and the correlation time *T*_*c*_ from the 50 samples using bootstrap methods [6]. To determine the error bars on 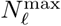, we use the distributions of 〈*V*_*x*_〉 generated from the bootstrap procedure. For fixed gradient and cluster properties, we draw a value of 〈*V*_*x*_〉 (*N*_*ℓ*_) for each possible value of *N*_*ℓ*_. Then, for each draw we record which value of *N*_*ℓ*_ gives the highest 〈*V*_*x*_〉. We draw 10,000 times to generate a distribution for 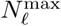. This procedure produces a distribution of the number of leaders which optimizes a given quantity.

We account for periodic boundary conditions for both the leader and the follower cells. For follower cells, we compute *θ*_*f*_ − *θ*_*c*_ with the arctan2 function as

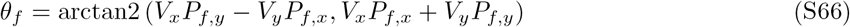

where the first term is the cross product **V**_*c*_ × **P**_*f*_ = |**V**_*c*_||**P**_*f*_ |sin(*θ*_*f*_ − *θ*_*c*_) and the second is the dot product **V**_*c*_ *·* **P**_*f*_ = |**V**_*c*_||**P**_*f*_ |cos(*θ*_*f*_ − *θ*_*c*_) and arctan2(b,a) returns the arc tangent of a vector 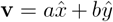. Therefore, Equation 8 will return *θ*_*f*_ − *θ*_*c*_, and the function is defined so that the angle is on the interval [−*π, π*]. For leader cells, we apply Equation S32 to their angles at each time step. This is equivalent to the procedure in Equation if *V*_*x*_ = 1 and *V*_*y*_ = 0—in either case, the angle relative to the x axis wrapped on [−*π, π*] is returned.

To avoid numerical errors associated with division by zero, the minimum of the concentration is taken as 10^−14^ to avoid dividing by zero when computing the leader cell uncertainty for cells far from the transition region. Since that is a value much lower than the dissociation constant, the leaders are just as random as they would be if the value were truly 0, and this does not impact the results.

**Figure S11:**
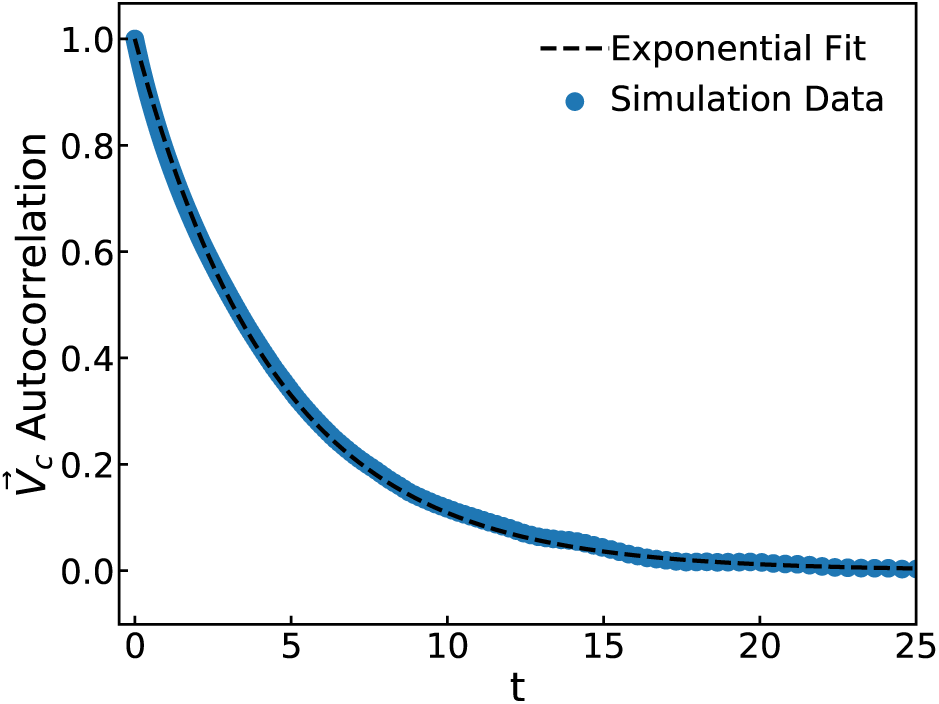
Representative exponential fit for autocorrelation function. The correlation function and an exponential fit are compared for a train of *N* = 50 cells in an *h* = 10 gradient with 10 leader cells and followers with Δ_*f*_ = 36°. We use the exponential fit to determine the correlation time.

